# Neural substrates for saccadic modulation of visual representations in mouse superior colliculus

**DOI:** 10.1101/2024.09.21.613770

**Authors:** Joshua B. Hunt, Anna Buteau, Spencer Hanson, Alon Poleg-Polsky, Gidon Felsen

## Abstract

How do sensory systems account for stimuli generated by natural behavior? We addressed this question by examining how an ethologically relevant class of saccades modulates visual representations in the mouse superior colliculus (SC), a key region for sensorimotor integration. We quantified saccadic modulation by recording SC responses to visual probes presented at stochastic saccade-probe latencies. Saccades significantly impacted population representations of the probes, with early enhancement that began prior to saccades and pronounced suppression for several hundred milliseconds following saccades, independent of units’ visual response properties or directional tuning. To determine the cause of saccadic modulation, we presented fictive saccades that simulated the visual experience during saccades without motor output. Some units exhibited similar modulation by fictive and real saccades, suggesting a sensory-driven origin of saccadic modulation, while others had dissimilar modulation, indicating a motor contribution. These findings advance our understanding of the neural basis of natural visual coding.

## INTRODUCTION

Animals actively explore their environment with orienting movements to acquire behaviorally relevant sensory information^1^. Interpreting sensory input, therefore, requires distinguishing between the sensory signals that arise from the environment and from the animal’s own movements. With respect to vision, decades of research under controlled conditions in the absence of self-generated motion has elucidated a hierarchical representation of visual information from the retina through higher-order brain regions^2–6^. However, how these representations are modulated by the movements that support ethological behavior is less well understood, preventing an understanding of natural visual processing.

Saccades, rapid eye-mediated gaze shifts, are common orienting movements that produce self-generated motion signals that must be taken into account by the visual system^7^. Saccades to visual targets decrease perceptual sensitivity to visual stimuli^8–11^ and suppress neural activity across the visual hierarchy^12–17^, phenomena commonly referred to as saccadic suppression. Target-directed saccades are just one of a diverse repertoire of eye movements observed during active vision in mammalian animal models such as primates^18^ and mice^19–23^. During natural visual exploration, the eyes and head are continuously brought out of alignment, eliciting frequent centripetal saccades to reset the eye-head angle to a default position. Such "resetting” saccades^21^ engage the same motion processing circuits as target-directed saccades and therefore present a similar challenge to visual coding^24^. Thus, determining how resetting saccades affect visual representations would advance our understanding of saccadic modulation more generally and is critical for understanding natural visual processing.

The superior colliculus (SC) contains neurons with diverse visual response properties^25,26^ critical for sensorimotor transformations^27–30^ and active visual functions^31–34^. The superficial layer of the SC receives direct retinal input^35^, which then influences motor processing in the intermediate and deep SC^36,37^. Suppression of neural activity by target-directed saccades was first described in the SC^13^, and slice studies suggest a simple underlying intracollicular mechanism by which superficial visual activity is inhibited by intermediate-layer neurons^38–40^. However, other studies have shown that signal processing in the retina itself can suppress responses during fast image translations^41–49^. Whether saccades modulate visual SC activity via sensory signals, motor programming, or both thus remains unclear.

We therefore sought to examine whether and how neural representations of visual stimuli are modulated by resetting saccades. We addressed these questions with an efficient and unbiased “white noise” approach that elicited frequent resetting saccades while recording responses of large populations of mouse SC neurons to frequent visual probes. We then compared visual responses in the absence of saccades and at a range of stochastically generated saccade-probe (S-P) latencies. We found that, across classes of functionally defined neurons, many exhibited saccadic suppression while some, surprisingly, exhibited enhancement, with the magnitude and sign of modulation dependent on the latency between the saccade and the visual response to the probe. By repeating these experiments under conditions that mimicked the visual experience during saccades, we found that saccadic modulation in some SC neurons required eye movement, while the modulation in others could be explained by the visual input alone. These findings suggest diverse and dynamic effects of saccades on visual representations and demonstrate multiple substrates for saccadic modulation in the SC.

## RESULTS

### Approach to examining saccadic modulation of visual representations in mice

To investigate how visual representations in the SC are modulated by saccades, we developed a paradigm to elicit temporally interpolated saccades and visually driven SC activity (Figure 1). Head-fixed mice in an immersive visual arena were presented with a low-contrast, full-field horizontally drifting sinusoidal grating (contrast=30%, spatial frequency=0.2 cycles/deg, velocity=12°/s; Figure 1B), which reliably elicits smooth tracking eye movements followed by resetting saccades (∼0.32 saccades/s), by engaging the optokinetic reflex^50^. The drift direction alternated between left and right for 90 s each, separated by 3-5 s of gray screen (Figure 1B). We recorded both pupils with high-speed cameras, estimated eye position by tracking the center of the pupil with DeepLabCut^51^ (Figure 1A) and extracted the initiation time and direction of saccades with machine learning (Methods). Figure 1C shows that leftward and rightward saccades are well-separated from each other and from non-saccadic eye movements. To elicit visual responses in the SC coincident with saccades, we also presented probe stimuli consisting of a transient increase in grating contrast (Figure 1B; duration=50 ms, contrast=100%; mean frequency=1.3 Hz), which yielded a sufficient range of S-P latencies for analysis (Figure 1D; negative latencies indicate that the probe precedes, and positive latencies that the probe follows, the saccade). Finally, to record SC visual activity, we targeted the superficial and intermediate layers of the SC with a Neuropixels electrode inserted tangential to the medial-lateral axis of the brain^52^ (26.9 ± 25 units/recording, mean ± standard deviation; Figures 1E, 1F, and S1; Methods). This experimental approach allowed us to examine how visual representations in a diverse population of SC neurons depend on the occurrence and timing of a coincident saccade (Figure 1G).

**Figure 1.**
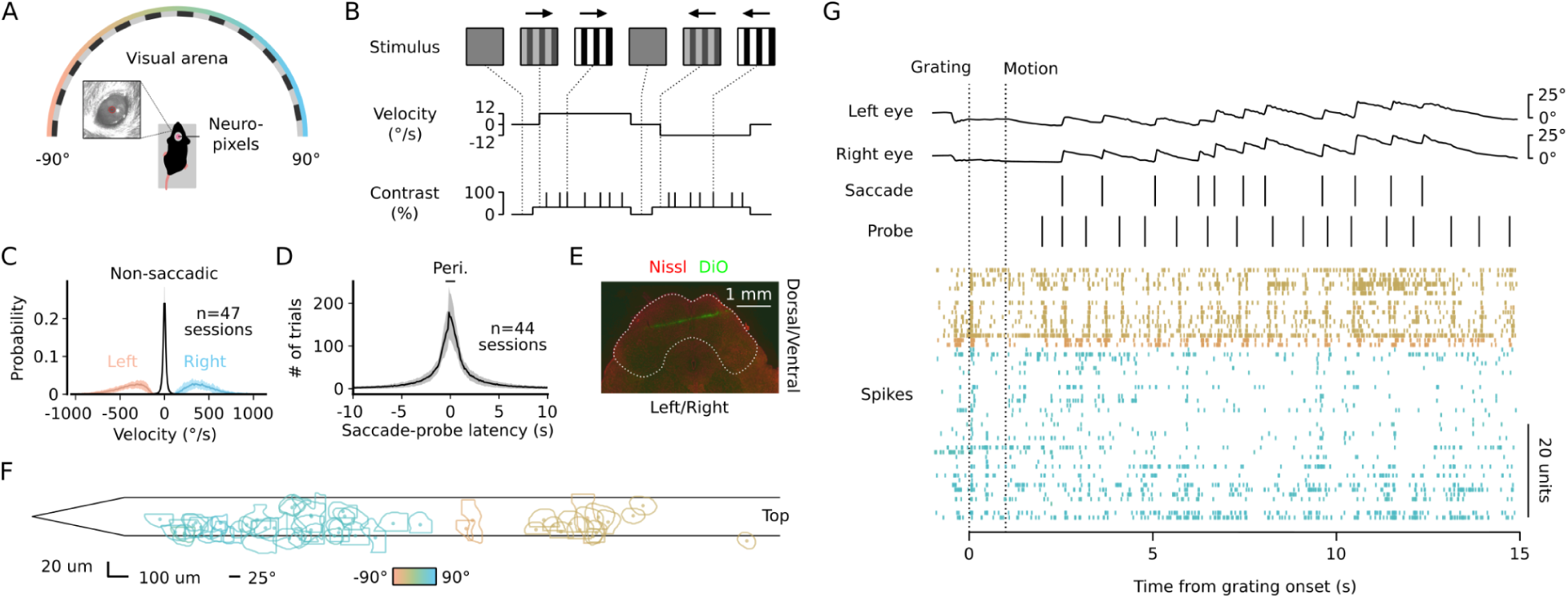
Strategy for studying saccadic modulation. **A.** Experimental design: head-fixed mice were presented with visual probes and drifting gratings to elicit saccades while cameras captured pupil positions and neuropixels probe recorded populations of visual SC units. **B.** Visual stimulus consisting of 90 s blocks of gratings drifting leftward or rightward with probes (0.05 s grating contrast increment) presented every 0.5-1 s. **C.** Distribution of peak eye movement velocity for leftward (orange) and rightward (blue) saccades or non-saccadic eye movements (black) across 47 sessions. Solid lines are means across sessions. Shaded region is mean ± 1 standard deviation. **D.** Distribution (mean ± 1 standard deviation) of S-P latencies for 44 sessions. Black bar indicates −0.5 to 0.5 s perisaccadic window used for subsequent analyses. **E.** Coronal SC section showing electrode tract (DiO, green); red, Nissl. Scale bar, ∼1000 µm. **F.** Receptive fields (RFs) for a subset of visual units from an example session, centered on each unit’s center-of-mass position along the electrode. Color indicates azimuthal position of the RF centroid. **G.** All experimental time series (eye position, saccades, probes and spikes) for a ∼15 s segment of the example session, units are color coded by their RF position shown in F. Dotted lines indicate the appearance of the grating stimulus (left) and then the onset of grating motion (right).

### Diverse visual representations in the SC are modulated by saccades

To determine how saccades modulate visual representations in the SC, we first characterized how the activity of visual SC units represents probes in the absence of saccades (‘extrasaccadic probes”). We recorded from 1,383 well-isolated single units in the superficial and intermediate layers of the SC with a detectable response to the probe (Figure S1; 44 sessions from 4 mice; ZETA test^53^, p<0.01; Methods). Because many units responded differently to probes drifting leftward and rightward, we focused on responses to probes in the preferred direction of each unit (Methods). We defined the response to extrasaccadic probes (R_Probe(Extra)_) as the standardized, trial-averaged peri-stimulus time histogram for trials in which the preferred-direction probe was presented without a saccade occurring within 500 ms. Consistent with previous work^25,26^, single units exhibited diverse temporal dynamics, varying in sign, complexity, and response latency (Figure 2).

**Figure 2.**
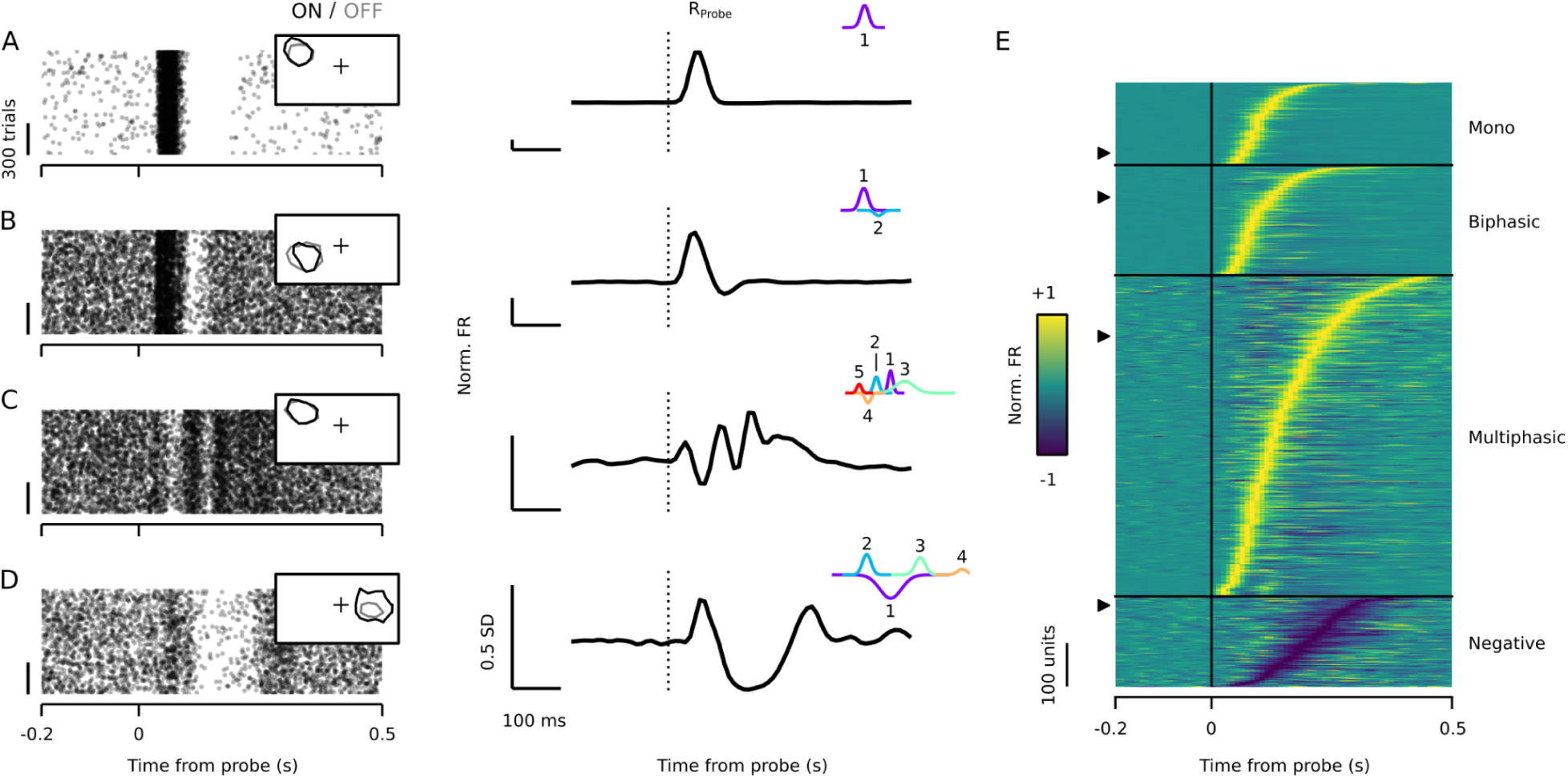
SC visual units exhibit diverse neural representations of probes. **A.** Raster (left) and trial-averaged, standardized peri-stimulus time histograms (PSTH; right) aligned to probe drifting in the preferred direction for an example unit with a positive, monophasic response. Left inset, ON (black) and OFF (gray) receptive fields. Right inset, individual components (shown in different colors) of the Gaussian mixtures model fit to the PSTH. Dashed line indicates the time of probe onset. **B-D**. Same as in A but for examples of biphasic, multiphasic and negative units, respectively. **E**. Amplitude-normalized PSTHs for all visual SC units used for subsequent analysis grouped by unit type. Arrows indicate examples in A-D.

To quantify this diversity, we approximated R_Probe(Extra)_ by fitting it with a Gaussian mixtures model (GMM; Methods). Some units exhibited a monophasic increase in firing rate best fit by a GMM with a single component (Figure 2A); whereas others exhibited a biphasic or multiphasic response that required two or more components to adequately fit (Figure 2B and 2C, respectively). The fourth class of units exhibited responses dominated by a decrease in firing rate and tended to have complex, multiphasic responses (Figure 2D). We next examined how saccades modulate responses to “perisaccadic probes” (R_Probe(Peri)_) by considering trials in which a saccade occurred within 500 ms of probe presentation. To account for any visual response to the saccade itself (R_Saccade_; Figure 3A), which often overlapped with perisaccadic probe responses, we first estimated R_Saccade(Shifted)_ by time-shifting R_Saccade_ by the latency between the probe and the saccade in each trial (Figure 3B) and averaging across trials (Figure 3C; Methods). We then isolated the perisaccadic probe responses (R_Probe(Peri)_) by subtracting R_Saccade(Shifted)_ from the combined response to the saccade and probe (R_Probe, Saccade_). Figure 3D and 3E shows R_Probe, Saccade_, R_Saccade(Shifted)_ and R_Probe(Peri)_ for trials in 100 ms bins of S-P latency (from −500 to 500 ms), for an example unit exhibiting a clear response to saccades, but a perisaccadic probe response unaffected by saccades (after accounting for the overlapping saccade response). Figures 3F and 3G show another example unit that responds to saccades, but exhibits a perisaccadic probe response that is almost entirely suppressed when the probe appears shortly after a saccade. Finally, we observed that some units lacked overt responses to saccades but nonetheless exhibited perisaccadic probe responses apparently modulated by saccades (Figures 3H and 3I). These examples demonstrate that R_Probe(Peri)_ can be isolated for units with and without saccade responses and suggest that saccades modulate visual representations in the SC of mice.

**Figure 3.**
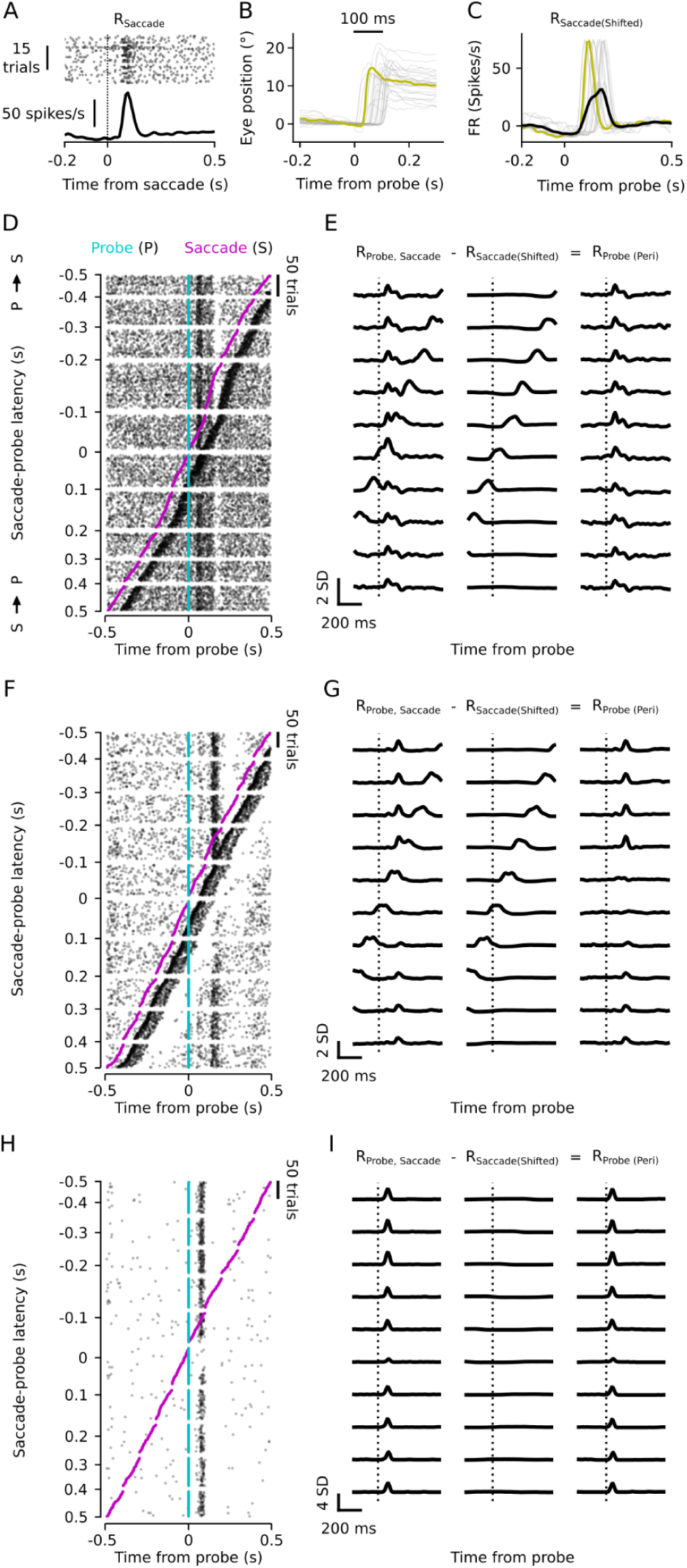
Saccades modulate responses to probes. **A.** Peri-saccade raster and trial-averaged firing rate (R_Sacccade_) for an example unit (same example as in D-E). **B.** Eye position as a function of time relative to probe onset for trials in which the saccade occurred 0 to 100 ms following probe presentation. Representative saccade waveform is highlighted in yellow, corresponding to the latency-shifted trace of R_Saccade_ in C. **C.** S-P latency-shifted traces of R_Saccade_ corresponding to trials shown in B in which the saccade begins 0 to 100 ms following probe onset. R_Saccade(Shifted)_, the average over all instances of R_Saccade_, is shown in black. The yellow trace corresponds to the example saccade in B. **D.** Raster for a representative unmodulated unit with trials sorted by S-P latency from −500 ms to 500 ms in 100 ms bins. Black ticks, spikes; cyan, probe; magenta, saccade. Negative/positive S-P latencies specify probes preceding/following the saccade, respectively. **E.** Approach for isolating probe responses on trials with coincident saccades. Left column, R_Probe, Saccade_; mean standardized firing rate for the observed perisaccadic response within each of the 100 ms time bins indicated in A. Middle column, R_Saccade(Shifted)_; the estimated saccade-related activity in each bin, calculated as in C. Note that the shape of R_Saccade(Shifted)_ depends on recorded saccade occurrences (magenta in D) and thus is distinct for each time bin. Right column, R_Probe(Peri)_; the difference between R_Probe, Saccade_ and R_Saccade(Shifted)_. The time of probe onset (cyan in D) is marked with a dashed vertical line. **F-G**. Same as in D-E, but for an example unit that exhibits a strong response to saccades and saccadic suppression. **H-I**. Similar to D-E, for an example unit that does not respond to saccades and exhibits saccadic suppression.

To quantify how saccades modulate responses to probes, we first compared R_Probe(Peri)_ to R_Probe(Extra)_ for each unit (Figure 4). Specifically, we compared the amplitude of the largest GMM component fit to R_Probe(Extra)_ to the amplitude of the same component from an identical model refit to R_Probe(Peri)_ (Figures 4A, 4B, and 4C; Methods). We then calculated the saccadic modulation index (MI) as the normalized difference between the amplitudes of the GMM fit to R_Probe(Peri)_ and R_Probe(Extra)_ such that MI < 0 indicates that saccades suppress probe responses, and MI > 0 indicates that saccades enhance probe responses (Figure 4D; MI significance estimated with bootstrapping, Methods). We found that a substantial fraction of units exhibited saccadic modulation, particularly at short S-P latencies. Most modulated units were suppressed by saccades (Figure 4D, blue), but some were enhanced (Figure 4D, red). Indeed, the frequency of modulation varied across time bins (X^2^=810.62 (18), p<0.001); suppression occurred more frequently when the probe followed the saccade (S→P), whereas enhancement most often occurred when the probe preceded the saccade (P→S, Figure 4E). We obtained similar results when we compared the extrasaccadic and perisaccadic probe responses of simultaneously recorded populations of SC units (Figures 4F, 4G, and 4H): the population vectors diverged during probe presentation (Figures 4F and 4G) and across sessions, this effect was strongest for short S-P latencies (n=44 populations, mean units/population=25.19; Figure 4H). These results demonstrate that visual responses in the SC are robustly modulated by the resetting saccades frequently performed during natural behavior.

**Figure 4.**
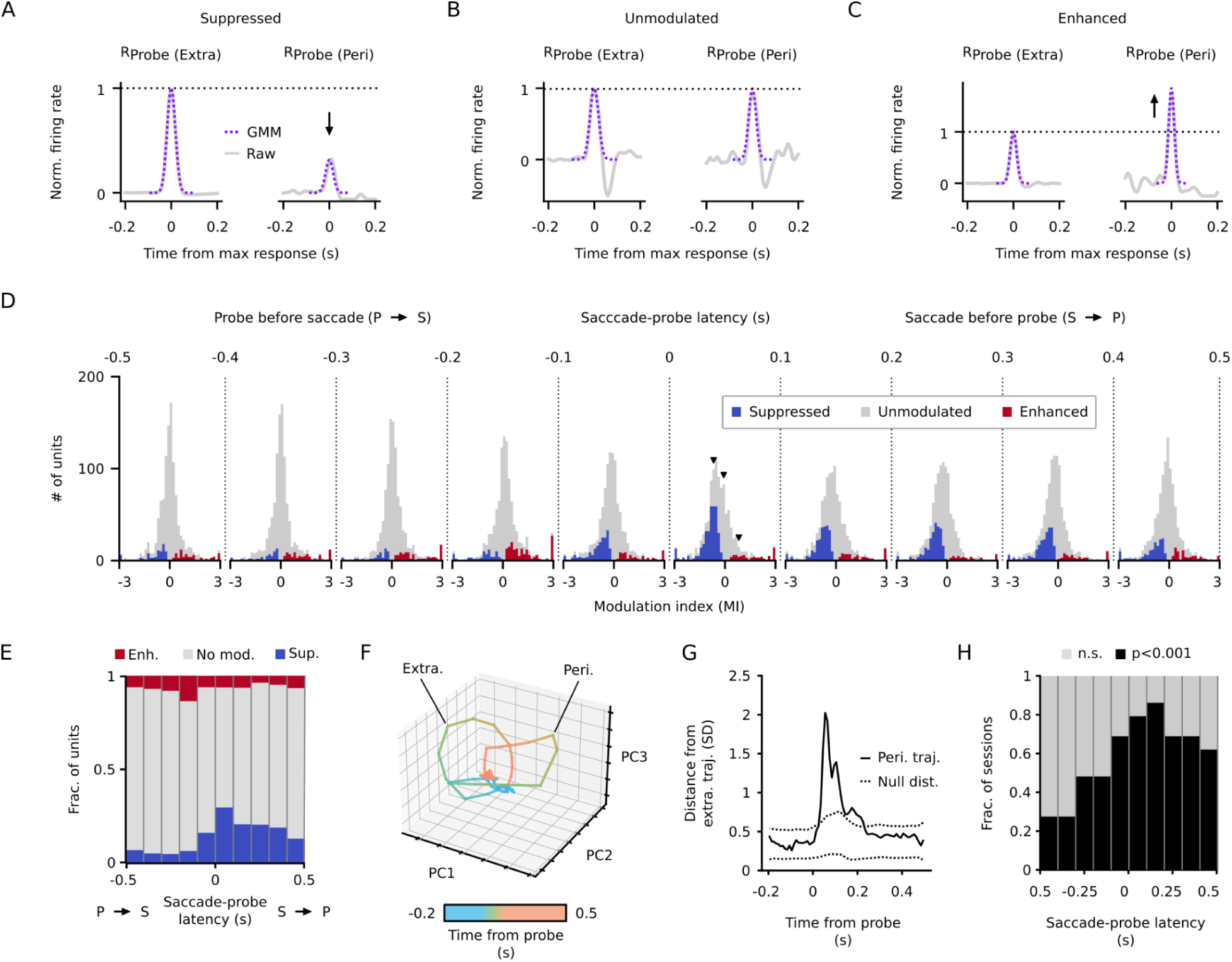
Saccadic suppression and enhancement exhibit distinct temporal dynamics. **A.** R_Probe(Extra)_ and R_Probe(Peri)_, left and right, respectively, for an example unit that exhibits saccadic suppression (S-P latency=0 to 100 ms). Dashed lines, GMM component with the largest amplitude. **B-C**. Same as in A but for a unit that is not modulated (B) or is enhanced (C). **D.** Saccadic modulation for all visual SC units (n=1383) as a function of S-P latency. Units with significant suppression or enhancement (p<0.05, bootstrapping test) are indicated in blue or red, respectively. Black arrows, examples from A-C. **E.** Fraction of units suppressed (blue), enhanced (red), or unmodulated (gray) for each 100 ms S-P latency bin. **F.** Neural trajectories representing the mean response to extrasaccadic probes (solid line) or perisaccadic probes (dashed line; S-P latency=0 to 100 ms) for an example population of simultaneously recorded visual SC units. **G.** Euclidean distance between extrasaccadic and perisaccadic population vectors for the example population shown in F. **H.** Fraction of sessions with a significant difference between extrasaccadic and perisaccadic population responses as a function of S-P latency.

### Saccadic modulation depends on the timing of saccades, probes, and visual responses

Consistent with previous studies performed in the primate SC^54–57^, our findings suggest that saccadic modulation is strongest at short S-P latencies (Figure 4 and Figure 5A). However, given that we observed modulation at a wide range of S-P latencies, we wondered if modulation might also depend on the latency between the probe and the maximum response (P-M latency; Figure 5A), or on the latency between the saccade and the maximum response (S-M latency; Figure 5A). Given that our dataset contains units with a range of P-M latencies (Figure 5B), and trials with a range of S-P latencies (Figure 1D), we can determine which latency best explains the temporal dynamics of saccadic modulation.

**Figure 5.**
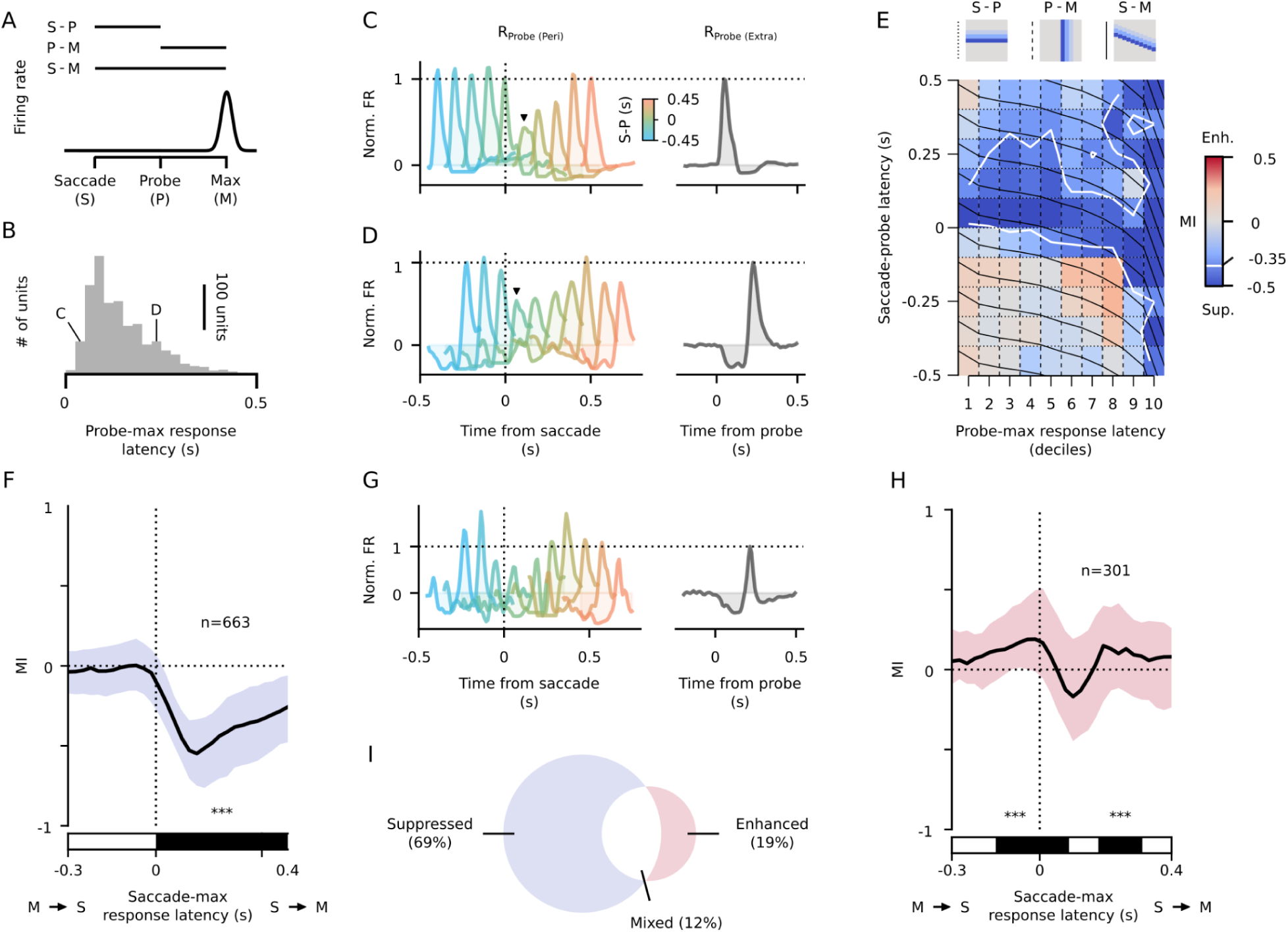
Modulation peaks at specific time windows around the saccades. **A.** Illustration of saccade-probe latency (S-P), probe-maximum response latency (P-M), and saccade-maximum response latency (S-M). **B.** Distribution of P-M latency for all visual SC units. **C.** Left, a family of R_Probe(Peri)_ responses for all S-P latency bins for an example unit with a short P-M latency. Right, R_Probe(Extra)_. Black arrow indicates R_Probe(Peri)_ with the lowest amplitude. Vertical and horizontal dotted lines indicate the time of saccade onset and the amplitude of R_Probe(Extra)_, respectively. Each individual R_Probe(Peri)_ shows the unit’s firing rate from 0 to 300 ms relative to probe presentation. **D.** Same as in C but for an example unit with a prolonged P-M latency. **E.** Top. Predicted pattern of suppression as a function of S-P and P-M latencies for three alternative hypotheses. Left, middle, and right cartoons indicate the pattern of suppression if suppression depends on S-P latency, P-M latency, or S-M latency, respectively. Lines to the left of the cartoons indicate the corresponding set of contours overlaid on the heatmap below. Bottom. Mean MI across all suppressed units (n=663) as a function of S-P latency (rows) and P-M latency (columns). Units on the left of the x-axis exhibit shorter P-M latencies whereas units on the right exhibit longer P-M latencies. This representation is constructed by first sorting the units based on their P-M latencies and assigning each unit to a single column. Then, the MI computed for each S-P latency is assigned to a row. Finally, the average MI is computed for each cell in this matrix. Dotted horizontal lines show expected contours if modulation depends only on S-P latency. Dashed vertical lines show expected contours if modulation depends only on P-M latency. Solid black lines show expected contours if modulation depends on S-M latency. White contours indicate mean MI < −0.35. **F.** Amplitude-weighted median MI for units exhibiting suppression but not enhancement for S-P latencies of −100 to 200 ms. Shading, interquartile range weighted by amplitude of R_Probe(Extra)_. Bottom bar shows latencies with median MI significantly different than zero (p<0.001, Wilcoxon signed-rank test). **G.** Same as in C but for a unit that exhibits enhancement. **H.** Same as in F but for units that exhibit enhancement during any S-P latency between −100 and 200 ms. **I.** The overlap of suppressed and enhanced units.

To illustrate this point, Figures 5C and 5D show R_Probe(Peri)_ for two example suppressed units that exhibit short (48 ms) and long (228 ms) P-M latencies, respectively. For each unit, we color-coded R_Probe(Peri)_ for trials in 100 ms bins of S-P latency (as in Figure 3) as a function of time relative to the saccade. This visualization allows us to examine each unit’s response dynamics relative to saccade initiation. If saccades modulate the representation of visual stimuli occurring at specific S-P latencies, we would expect to observe the strongest modulation at the same S-P latency, indicated by the same color of R_Probe(Peri)_, for the two units. However, the strongest modulation was instead observed when each unit’s maximum response occurred around 100 ms after the saccade (Figure 5C and 5D, black arrows), regardless of the S-P latency, suggesting that modulation depends primarily on S-M latency.

We quantified this analysis across our population of units exhibiting significant saccadic suppression in any 100 ms window of S-P latencies from −100 to 200 ms (n=663 units) by binning the MI within units by S-P latency (Figure 5E, bottom, rows) and across units by their P-M latency (Figure 5E, bottom, columns). If the strength of suppression depends only on S-P latency, MI will vary only across the rows of this matrix (Figure 5E, top, left); if it depends only on P-M latency, MI should vary only across the columns (Figure 5E, top, middle); and if it depends on S-M latency, MI would vary along both rows and columns (Figure 5E, top, right). These predictions are illustrated by the cartoons at the top of Figure 5E and by the black contour lines superimposed on the matrix in the bottom of Figure 5E. The white contour indicates the mass of saccadic suppression in this 2-dimensional latency space, which closely matches the predicted results were modulation to depend on S-M latency (Figure 5E, bottom, solid black lines). Indeed, plotting median MI as a function of S-M latency for these suppressed units reveals that suppression is maximal when the response maximum occurs at a similar latency – ∼100 ms – after the saccade (Figure 5F).

We performed the same analysis with units that exhibited significant saccadic enhancement in any 100 ms window of S-P latencies from −100 to 200 ms (n=301 units). R_Probe(Peri)_ binned by S-P latency (as in Figures 5C and 5D) for an example enhanced unit is shown in Figure 5G. Interestingly, saccadic enhancement appears to be multiphasic: this example unit is enhanced when the response maximum precedes the saccade, suppressed when the response maximum shortly follows the saccade, and again enhanced at longer S-M latencies. This pattern was consistent across enhanced units. Notably, for some units, enhancement precedes saccade initiation (Figure 5H). In line with our finding that many enhanced units also exhibit suppression around 100 ms after saccade initiation (Figure 5H), we found that 12% of modulated units exhibited both suppression and enhancement, at different S-M latencies (Figure 5I).

### Saccadic modulation is independent of directional tuning

Given that the probes were presented during either leftward or rightward grating motion, many visual neurons in mouse SC are direction selective^58^, and OKR is tightly linked to direction selectivity circuitry^24^, we investigated the relationship between saccadic modulation and direction selectivity. We first computed a horizontal direction selectivity index (hDSI) for each unit as the absolute value of the difference between the average responses to leftward and rightward drifting gratings normalized by their sum, which quantifies the strength of directional preference agnostic of the preferred direction. Across the population, we observed a range of directional tuning for visual stimuli, from strong (Figure 6A) to weak (Figure 6B), and a similar preference for horizontal motion direction when assessed with other stimulus ensembles and saccades (Figure S2). Using a threshold of hDSI >= 0.3, 13.1% of units were identified as being direction selective (DS; Figure 6C). DS cells largely mirror the saccadic modulation patterns observed for non-DS cells, exhibiting nearly identical frequencies of saccadic modulation as a function of S-P latency (Figure 6D). Additionally, we observed a strong positive correlation between MI in the preferred and null directions for both DS (r=0.22, p=0.013) and non-DS cells (r=0.78, p<0.001; Figure 6E). Thus, saccadic modulation appears to operate independently of directional tuning.

**Figure 6.**
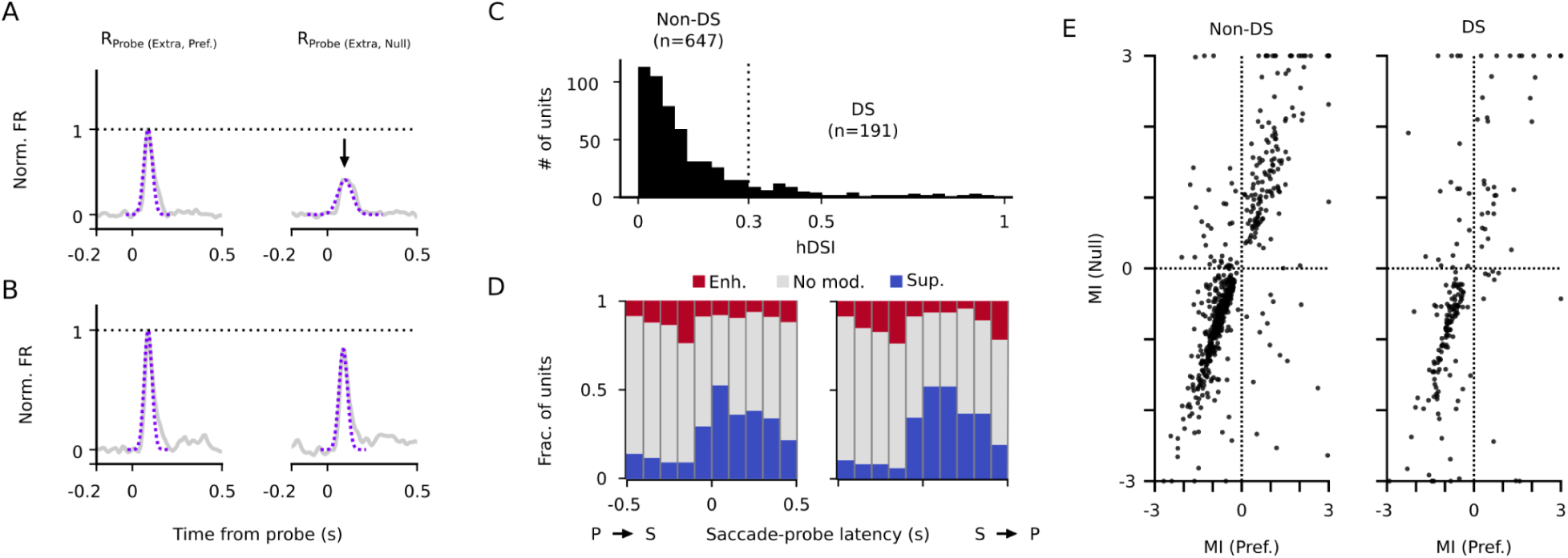
Saccadic modulation does not depend on directional tuning. **A.** R_Probe(Extra)_ for probes in the preferred or null direction for an example direction-selective unit. **B.** Same as in A for an example unit that is not direction-selective. **C.** Distribution of horizontal direction-selectivity indices across units. Units with low signal-to-noise ratios (max(abs(R_Probe(Extra, Pref.)_)) < 0.5 SD) were excluded from this analysis. **D.** Fraction of units that are enhanced (red), unmodulated (gray), or suppressed (blue) as a function of S-P latency for non-DS units (left) and DS units (right). **E.** Correlation between MI calculated for probes in the preferred and null directions, for non-DS and DS units.

### Visual and motor signals contribute to saccadic modulation

The fact that some units exhibit saccadic modulation when their probe response precedes the saccade (Figure 5H) suggests a non-visual mechanism of modulation, consistent with the idea of a saccade-related corollary discharge signal modulating visual activity^39,57,59^. However, other findings demonstrate that the visual signal experienced during the saccade can itself modulate visual representations as early as the retina^41,42^. To address the contributions of visual and motor mechanisms to the saccadic modulation we observed in visual SC neurons, in a subset of sessions we compared, within units, saccadic modulation produced by fictive saccades - rapid stimulus translations designed to mimic the visual experience of the saccade - to modulation produced by real saccades (Figure 7A; Methods).

**Figure 7.**
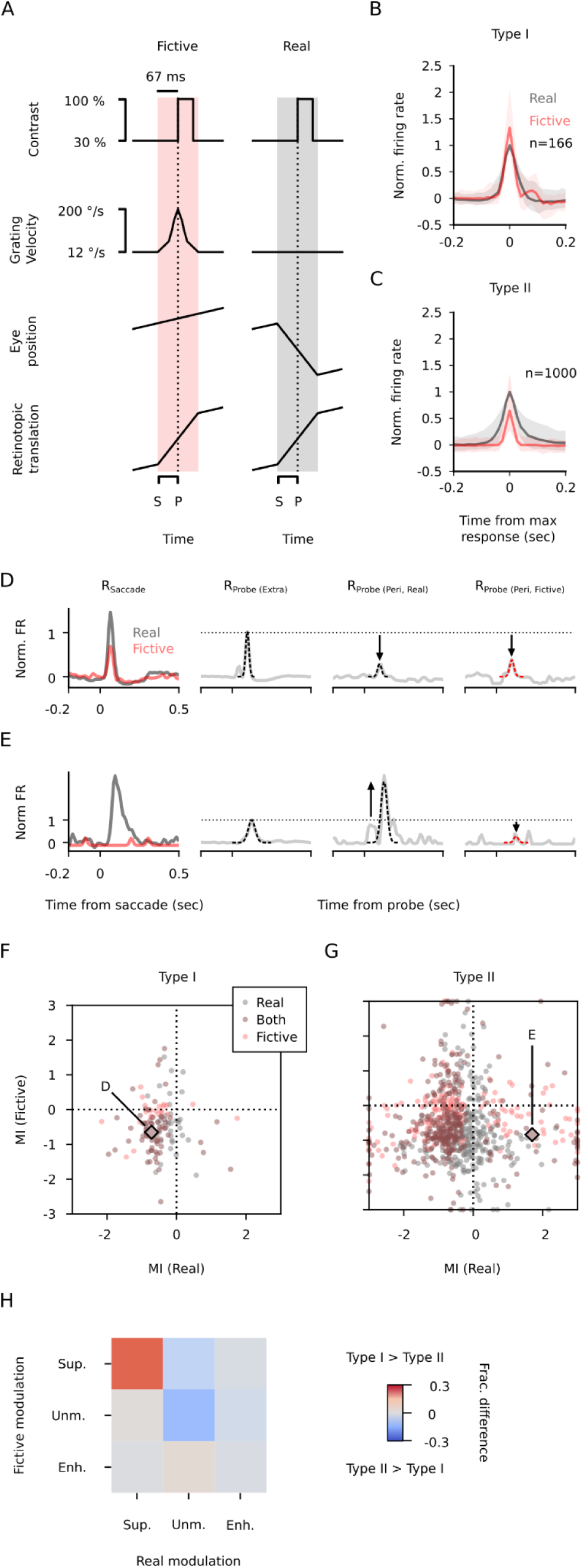
Saccadic modulation arises from visual and non-visual mechanisms. **A.** Schematic showing how fictive saccades (left) mimic the retinotopic translation of the probe (bottom) during real saccades (right). Saccade (S) and probe (P) times are indicated on the bottom. **B.** Median (±1 quartile) firing rate during real (gray) and fictive (red) saccades (R_Saccade_) for units exhibiting a strong response to both (Type I). Firing rate is normalized to the amplitude of the response to real saccades and centered on the maximum of the response. **C.** Same as in B but for units that exhibit a weak response to fictive saccades (Type II). **D.** Example Type I unit that exhibits saccadic suppression to both real and fictive saccades. From left to right, subplots show the response to saccades, and probes presented near real and fictive saccades. **E.** Same as in D but for an example Type II unit that is enhanced by real saccades and suppressed by fictive saccades. **F.** The modulation of most Type I units is similar between real and fictive saccades. Units are color coded based on the significance of their modulation. Units not significantly modulated by real or fictive saccades are not shown. **G.** Same as in F but for Type II units. **H.** Difference of the joint distribution of saccadic modulation for Type I and Type II units. Red indicates that a higher fraction of Type I units was observed than Type II units; blue indicates that a higher fraction of Type II units was observed than Type I units.

We first found that some units responded to real and fictive saccades equivalently (Type I, 166/1166 units) while others did not respond to at least one type of saccade (Type II, 1000/1166 units), and there was a modest association between these Types and the complexity of extrasaccadic responses (X^2^ (3), p=0.02). Specifically, Type I units were more frequently biphasic (+9%; Figure 2B), and Type II units were more frequently monophasic (+3%; Figure 2A), in their responses to extrasaccadic probes. The robust responses of Type I units to real and fictive saccades suggests that the fictive saccade paradigm succeeded in mimicking the visual stimulus generated by real saccades (Figure 7B). Most Type II units had weak responses to fictive saccades, suggesting that their responses to real saccades may reflect the input of a saccade-related motor signal.

We calculated R_Probe(Peri, Real)_ and R_Probe(Peri, Fictive)_ as the perisaccadic responses for real and fictive saccades, separately, and used these terms to calculate MI_Real_ and MI_Fictive_ for each unit (Figures 7D and 7E). In these experiments, fictive S-P latency was always within 0 to 100 ms, so we used the same real S-P latency range for comparison. For Type I units, we found that the joint distribution of real and saccadic modulation was nonuniform, with most units exhibiting suppression by both real and fictive saccades (42%; X^2^=199.2 (8), p<0.001; Figure 7F), consistent with a visual mechanism for saccadic suppression. For Type II units, while fictive saccades elicited weak responses on their own, they frequently suppressed probe responses (48% of Type II units suppressed), similar to the suppression by fictive saccades in Type I units (63% of Type I units suppressed). However, Type II units exhibited modulation by real saccades distinct from Type I units, characterized by less frequent suppression (60% to 38%) and more frequent enhancement and absence of modulation (4% to 9% and 36% to 52%, respectively; X^2^=5211.7 (2), p<0.001; Figures 7G and 7H). Together, these results suggest that a motor signal mediating saccadic modulation may counteract the suppression arising from visual signaling, demonstrating that saccades modulate visual representations through both motor and visual mechanisms.

## DISCUSSION

We examined how resetting saccades modulate visual representations in the SC, a critical locus for coordinating motor output in response to visual input. We found that, despite the diversity in how SC units responded to a simple contrast probe (Figure 2), and regardless of how selective the unit was for the direction of probe motion (Figure 6), many SC units exhibited modulation (Figures 3-7). The majority of modulated units were suppressed, although a significant fraction exhibited enhancement (Figures 4C and 4E; 5G-I). Further, we found that the timing and sign of saccadic modulation were best explained by the latency between the saccade and the neural response, rather than between the saccade and the probe itself (Figure 5). Finally, we found that the visual signal alone was sufficient to account for saccadic modulation in a subset of units, while in other units, the eye movement itself plays a role (Figure 7). These results demonstrate that visual representations in the SC are modulated by resetting saccades, which could contribute to accurate visuomotor transformations during behavior.

Naturally behaving animals make orienting movements to obtain behaviorally relevant information about their environment^7,22^. In primates, the angle between the head and eyes frequently deviates from its default alignment^60–62^, which elicits centripetal resetting saccades that recover the default angle of the eyes^63–65^. During active vision in mice, resetting saccades similarly return the eyes to a default angle as part of head-initiated gaze-shifts^19,20,23^. These studies demonstrate that resetting saccades present a frequent challenge to the visual system across mammalian species. By using the OKR to elicit centripetal saccades that return the eyes to their default angle within the head, we show that resetting saccades, despite serving a distinct function than target-directed saccades, also modulate visual representations.

The modulation of visual responses that we observed with resetting saccades broadly resembles previous findings with target-directed saccades. Foundational studies in primates identified a subset of SC neurons that responded differently when a stimulus translated across their receptive fields than to the same stimulus traversed by the receptive field during a saccade^13^. Subsequent research built upon this finding by demonstrating that visual responses in the SC are attenuated around the time of microsaccades^54–56^ and target-directed saccades^57^. Our results complement this body of research, demonstrating that saccadic modulation generalizes to resetting saccades and to the mouse, despite fundamental differences between primates and mice in how saccades support visual behavior^21^.

Our findings also significantly advance our understanding of saccadic modulation in the SC. First, previous studies suggested that the magnitude of saccadic modulation depends on the S-P latency^54–57^. By recording from SC units exhibiting a large range of response latencies (Figure 2) at stochastically-generated S-P latencies (Figure 1D), we disambiguated the timing of saccades, probes and neural responses and found that saccadic modulation depends on the latency between the saccade and the maximum response to the probe (Figure 5). In particular, we found that suppression of responses to probes presented prior to the saccade (Figures 4D, 4E and 4H) could be better accounted for by considering the timing of the response itself, which typically followed the saccade (Figure 5F). In contrast, enhancement of probe responses often occurred prior to the saccade (Figure 5H). These findings may have implications for how specific types of visual SC neurons, which exhibit different latencies in response to visual stimuli^25^, represent the visual scene during natural behavior. Future studies can determine how genetically-identified classes of visual SC neurons are modulated by saccades by targeting them for recording during our paradigm.

Second, our fictive saccade experiments (Figure 7) reconcile findings demonstrating that saccadic modulation can be explained by visual signals alone with other work supporting a role for motor signals. Previous studies suggest that saccadic suppression in the SC and other regions may be inherited from motion processing in the retina, which evolved to efficiently represent the dynamics of the visual scene during natural behavior and contains many neuron types that respond robustly to rapid global translation^41–49^. In particular, recent work posits that saccade-induced retinal image shifts facilitate suppression through signal filtering by photoreceptors^42^, and motion processing is further diversified by retinal amacrine cells that can inhibit retinal ganglion cell activation during fast motion^45,46,48,66–68^. Consistent with these findings, we found that probe responses in SC units with robust responses to fictive saccades are generally suppressed (Figure 7F).

However, given the diverse saccadic modulation exhibited by units with weak responses to fictive saccades (Figure 7G), we speculate that a motor signal complements the sensory drive to fine-tune the modulation of visual representations in the SC. Indeed, studies in rodent SC slices support a circuit mechanism linking a corollary discharge signal from saccade-initiating SC neurons to suppression of visual SC representations via inhibitory SC neurons^38–40,57^. Alternatively, given the presence of a parallel excitatory connection between premotor neurons and putative narrow-field cells in the superficial layers of the SC^69^, the motor signal may contribute specifically to the enhancement of visual representations^41^. In support of this hypothesis, we identified a significant subset of units that exhibited saccadic enhancement both before and after saccades. This finding is consistent with a previous report of saccadic enhancement of visual responses in the SC prior to microsaccades^55^. Such enhancement may contribute to perceptual advantages afforded by saccades, such as increased spatial attention to saccade targets^70^.

In addition to saccadic modulation, a potential effect of the motor signal is to change the directional tuning during a saccade, as observed in mouse V1^71^. However, our analysis of the representation of visual probes as well as fictive and real saccades indicates that directional preference of visual SC neurons does not change during the saccade (Figures 6 and S2). Similarly, directional tuning is conserved in motor SC neurons^37^, suggesting that saccades differentially affect motion processing in SC and cortex.

This study advances our understanding of the neural basis of vision during natural behavior. Future studies can interrogate the circuit mechanisms of saccadic modulation by recording and perturbing its key elements, such as inhibitory SC neurons known to mediate orienting movements^72^, examine how orienting head and trunk movements frequently observed during natural behavior modulate visual representations in the SC, and determine how the SC interacts with other brain regions to mediate active vision.

## ACKNOWLEDGMENTS

We thank Drs. Baohua Liu, Jamie Mazer, Daniel Denman, Ethan Hughes, Abigail Person, Diego Restrepo, and Cristin Welle as well as members of the Felsen and Poleg-Polsky labs for their insightful comments on the manuscript. We thank Dr. Michael Hall for machining, Dr. Ryan Williamson for engineering support, and Andrew Scallon for engineering support. Light microscopy was performed at the University of Colorado Anschutz Medical Campus Advanced Light Microscopy Core and engineering support was provided by the University of Colorado Optogenetics and Neural Engineering Core and The Innovation and Design for Experimentation and Analysis Core. All cores are supported in part by the Rocky Mountain Neurological Disorders Center (P30NS048154), by NIH/NCRR Colorado CTSI grant UL1 RR025780, and by the University of Colorado NeuroTechnology Center. This work was supported by the National Institutes of Health (R01EY035293, R01NS079518, R01NS129608, F31EY033651).

## AUTHOR CONTRIBUTIONS

Conceptualization, J.B.H, G.F., and A.P.-P.; Software, J.B.H. and S.H.; Formal analysis, J.B.H. and S.H.; Investigation, J.B.H. and A.B.; Data curation, J.B.H. and A.B.; Writing - Original draft, J.B.H., G.F., and A.P.-P.; Writing - Review and editing, J.B.H., G.F., and A.P.-P.; Funding acquisition, J.B.H., G.F., and A.P.-P., Supervision, G.F. and A.P.-P.

---

**Supplementary Figure 1.**
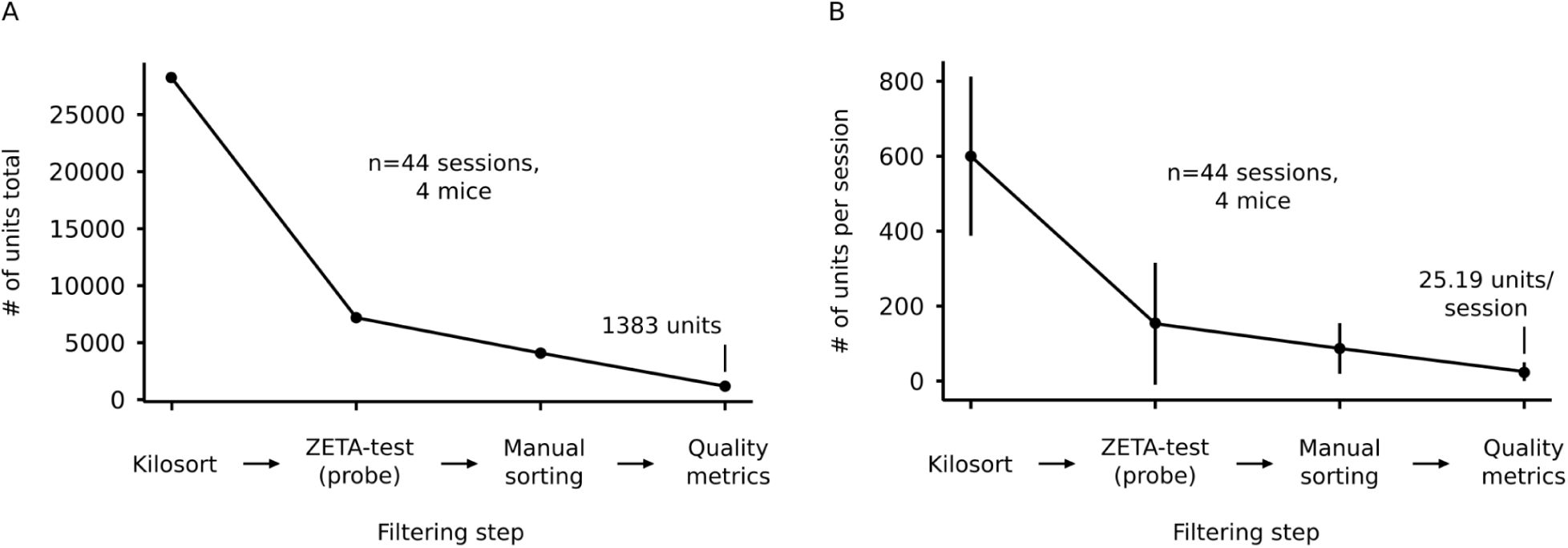
Processing stages for isolating single-unit activity from high-density extracellular recordings. **A.** Total number of units across 44 recordings and 4 animals at each step of the unit processing pipeline. **B.** Same as in A but showing the average number of units per recording ± 1 standard deviation.

**Supplementary Figure 2.**
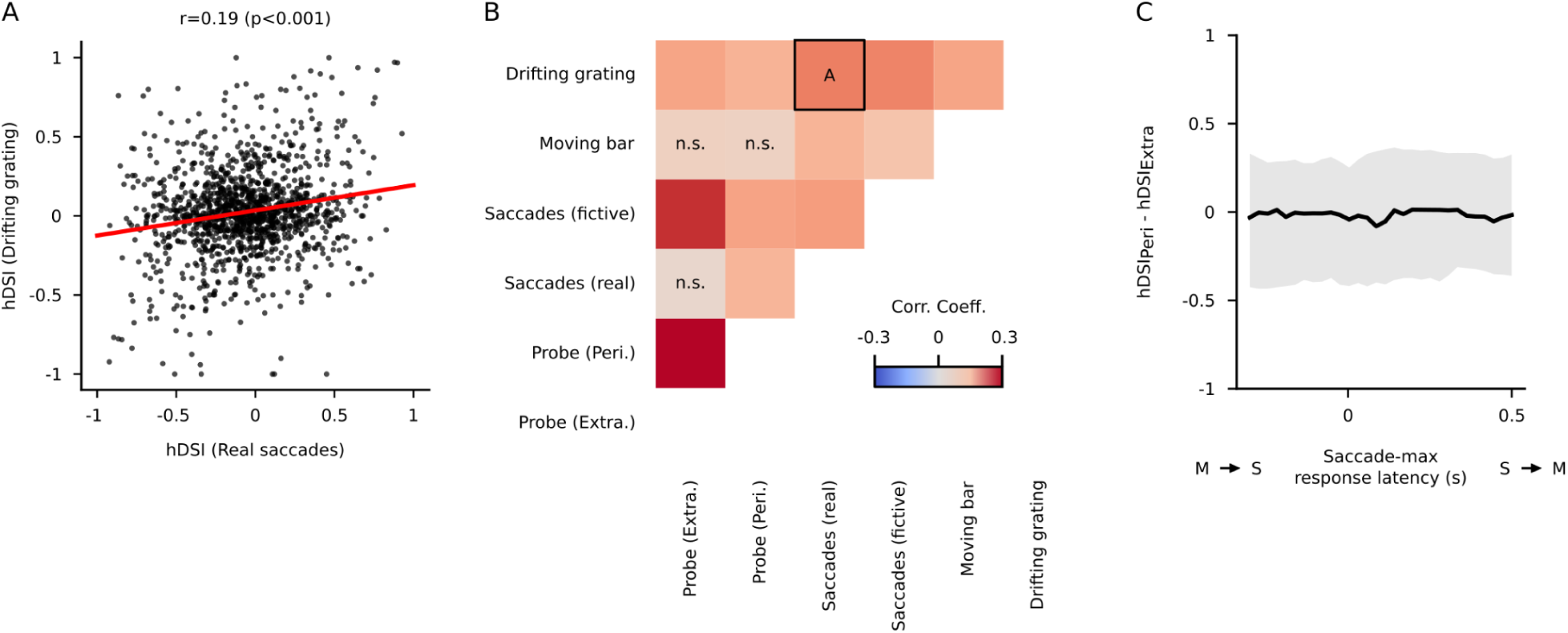
Direction preference is stable around the time of saccades. **A.** Correlation between signed hDSI for real saccades and signed hDSI for the drifting grating stimulus. Negative hDSI values indicate preference for leftward motion and vice versa. **B.** Correlation coefficients for correlations of signed hDSI between all stimuli with directionality. The scatterplot from A is indicated with a black square. **C.** Median difference between perisaccadic hDSI and extrasaccadic hDSI based on probe responses for units with an hDSI ≥ 0.3 as a function of saccade-maximum response latency. Shaded region indicates interquartile range.

## STAR METHODS

### Key resources table

| REAGENT OR RESOURCE | SOURCE | IDENTIFIER |
| --- | --- | --- |
| Experimental models: Organisms/strains |  |  |
| Mouse: Gad2-Cre | Jackson | RRID: ISMR_JAX:010802 |
| Software and algorithms |  |  |
| Python 3.9.13 | www.python.org/ | RRID: SCR_008394 |
| PsychoPy | www.psychopy.org/ | RRID: SCR_006571 |
| DeepLabCut | github.com/DeepLabCut/ | RRID: SCR_021391 |
| Open Ephys | open-ephys.org/gui | RRID:SCR_021624 |
| Kilosort 2.0, 2.5, 3.0 | github.com/MouseLand/Kilosort | RRID:SCR_016422 |
| Phy | github.com/cortex-lab/phy | N/A |

## RESOURCE AVAILABILITY

### Lead contact

Further information and requests for resources should be directed to and will be fulfilled by the lead contact, Gidon Felsen.

### Materials availability

This study did not generate new unique reagents.

### Data and code availability

Electrophysiological data reported in this paper will be shared by the lead contact upon request. All original code has been deposited at Zenodo and is publicly available as of the date of publication. DOIs are listed in the key resources table. Any additional information required to reanalyze the data reported in this paper is available from the lead contact upon request.

## EXPERIMENTAL MODEL AND SUBJECT DETAILS

All experiments were conducted in accordance with institutional guidelines with methods approved and overseen by the University of Colorado IACUC. All mice were housed with standard care in an accredited veterinary facility with free access to food and water.

### Animals

These experiments were performed on Gad2-cre mice (Jackson Labs #010802).

## METHOD DETAILS

### Visual arena

Animals were head-fixed and habituated to a low-friction rodent-driven belt treadmill^73^ for approximately one week preceding experiments. For visual stimulation, mice were placed within an immersive visual arena consisting of an LED projector (LightCrafter 3010 EVM), a spherical mirror (Edmund Optics), and an acrylic dome painted with projector screen paint (Screen Goo 2.0). The visual arena covers approximately 180 degrees of azimuth and 110 degrees of elevation (Figure 1A). Images projected onto the interior of the dome were transformed using open-source image-warping software for projecting images onto non-planar surfaces (PsychoPy).

### Eye tracking and saccade detection

Video recordings of eye movements were obtained with two FLIR USB3 monochrome cameras (one for each eye) acquiring at 200 frames/s. The position of the pupil center was tracked offline using open source pose estimation software (DeepLabCut). Saccadic eye movements were mostly conjugate (Figure 1H), so to simplify our data processing pipeline, we only considered eye position data from a single eye (left). To identify saccades, we first extracted high-velocity eye movements. For a given recording, any eye movement with a velocity greater than or equal to the 99th percentile of all eye velocities was considered a putative saccade. Putative saccade velocity waveforms (± 200 ms relative to peak velocity) were fed into a chain of machine learning models. The first model, a multi-layer perceptron classifier (scikit-learn), classified putative saccades as nasal, temporal, or non-saccadic eye movements. The subsequent model, a multi-layer perceptron regressor (scikit-learn), estimated the start and stop times of saccadic eye movements output by the first model. Both models were trained using manually collected training data from multiple animals across multiple sessions to prevent overfitting to a single animal or session.

### Visual stimuli

Mice were presented with drifting gratings to elicit saccades via the optokinetic reflex, sparse noise to map receptive fields, and fictive saccades to mimic the visual experience during saccades.

#### Sparse noise

We used sparse noise^74^ to estimate the receptive fields of visual units in the SC. Starting with minimal background luminance for each 10° x 10° subregion in a 10 x 17 grid centered on the visual arena, we increased the luminance of the subregion from black to white, then decreased luminance from white to black. Each phase of this ON-OFF sequence lasted approximately 500 ms, and there was no delay between the end of the OFF phase and the beginning of the next ON phase. The order of subregion activation was pseudo-randomized such that each subregion cycled through the ON-OFF sequence three times but not consecutively.

#### Drifting grating

To elicit the optokinetic reflex, mice were presented with a drifting grating stimulus. We selected parameters for the drifting grating that are optimal for eliciting the OKR in mice^50^: spatial frequency=0.2 cycles/degree and velocity=12 degrees/s. Sessions were organized into blocks of 60 or 90 s during which the grating appeared, remained static for a 1 s delay, then began to rotate clockwise or counter-clockwise. The sequence of blocks was pseudo-random such that motion direction (clockwise vs. counter-clockwise) was equally represented but unpredictable. The baseline contrast of the grating was set to 30%. During the motion of the grating, every 0.5 to 1 s, we incremented the contrast to 100% for 50 ms. The animals were shown a gray screen (contrast=0%) for 5 s in between blocks.

#### Fictive saccades

To simulate the visual experience of the drifting grating stimulus during real saccades, every 1-2 s the velocity of the grating was increased from 12°/s to 200°/s for 117 ms using a Gaussian function (µ=67 ms, σ=2.5 ms) to shape the velocity profile (Figure 7A). On 1/3 of trials, halfway through the fictive saccade (∼67 ms after the onset of the fictive saccade), we presented the probe stimulus (grating contrast increment from 30% to 100%; Figure 7A). The other two thirds of trials consisted of either the fictive saccade or the probe stimulus presented in isolation. For analysis (Figure 7), we excluded any trial with a real saccade that began within 100 ms of the fictive saccade or probe.

### Extracellular recordings

Extracellular neuronal recordings from the superficial and intermediate layers of the caudal SC were collected with Neuropixels 1.0 electrodes. Neural data acquisition was controlled with open-source software (Open Ephys). On the day before the first recording, a craniotomy (and durotomy) ∼3 mm in diameter was made immediately above the electrode insertion target. Craniotomies were made at 0.15 ± 0.13 mm (relative to lambda) along the anterior-posterior axis and 2.38 ± 0.22 mm along the medial-lateral axis. Immediately before recordings, animals were lightly anesthetized with isoflurane (1-2%), and the craniotomies were cleared of tissue regrowth. Using a stereotaxic micromanipulator (Kopf), the electrode was inserted into the center of the craniotomy, tangential to the medial-lateral axis of the brain at an angle of 14.3 ± 6.4 degrees and lowered to a depth of 1.7 ± 1.7 mm from the surface of the brain. We waited 15-30 minutes after inserting the electrode before starting the recordings to minimize electrode drift. Between recordings, the craniotomies were sealed with silicone (Kwikseal). On the day of the final recording for each animal, the electrode was coated in fluorescent dye, DiI or DiO (Invitrogen), immediately before insertion. Electrode placement within the SC was confirmed histologically (Figure 1F).

### Spike sorting and unit filtering

We used Kilosort 2.0, 2.5, and 3.0 to automatically sort extracellular spikes into single- and multi-unit clusters. We then used the ZETA-test^53^ to identify units with significant probe or saccade-related activity (p<0.01). We used open-source spike-sorting software (Phy) to manually curate the automatic spike-sorting results, focusing on merging partial single units judged to have been erroneously split by Kilosort and ensuring quality control by recategorizing single-units as noise or multi-unit activity based on firing rate over the entire recording, spike waveform shape, and inter-spike interval distributions. Finally, we discarded any unit that did not meet all threshold levels of quality for four commonly used spike-sorting quality metrics^6^: firing rate (≥ 0.2 Hz), presence ratio (≥ 0.9), amplitude cutoff (≤ 0.1), and inter-spike interval violation rate (≤ 0.1).

### Gaussian mixtures model

For each unit, we created a template of the extrasaccadic probe response which indicates how each unit represents the probe outside of the influence of saccades (R_Probe(Extra)_). First, we computed a peri-event time histogram (PETH) which quantified the firing rate in 10 ms time bins from −0.2 s to 0.5 s relative to extrasaccadic probes (distance to nearest saccade > 0.5 s). Next, the baseline level of firing rate in the window from –0.2 to 0 s was subtracted from each PETH. Finally, the PETH was scaled by the standard deviation of firing rate in 10 ms time bins sampled from across the entire recording. We approximated the extrasaccadic probe response by fitting R_Probe(Extra)_ with a Gaussian mixtures model (GMM; Figure 2). For each unit, we identified k unsigned peaks in R_P(Extra),_ where k is an integer between 1 and 5. If >5 peaks were detected, we only considered the 5 largest amplitude peaks. We then seeded a k-component GMM with an initial parameter estimate based on the amplitude, latency, and width of each detected peak. This model was then fit to R_Probe(Extra)_ using a least squares method for optimization. During the curve fitting, we constrained the GMM such that the amplitude of each peak could only vary ± 0.01 standard deviations from the initial estimate of amplitude, the peak latency could only vary ± −3 ms from the initial estimate of latency, and the peak width was within the range 1-20 ms. We constrained the curve-fitting in this way to minimize the overlap in individual components while still allowing us to extract amplitude and latency parameters for each component. With this approach, the GMM functions to decompose the response into distinct components instead of simply finding the best approximation of the response.

We chose to use a GMM to approximate the probe responses over a simpler method for measuring amplitude for several reasons. First, given that we frequently observe multiphasic responses (Figure 2), we reasoned that individual components of these responses could be differentially modulated by saccades. If we used an integrative metric to measure response amplitude, differential modulation of individual components might cancel out or dilute our estimate of modulation. Using a GMM to decompose multiphasic responses into isolated components allowed us to avoid this potential issue by considering modulation on a component-by-component basis. Second, the GMM functions as a dynamic response window such that measurements of response amplitude don’t depend on the temporal dynamics of the response. By using the GMM, we avoided having to choose the location and size of a static response window for each unit. Finally, we chose to use the GMM because it is robust to noisy signals. Given that some units, particularly the multiphasic units, exhibited relatively low signal-to-noise ratios, we reasoned that a simpler method for measuring response amplitudes (e.g., peak-finding) might result in imprecise estimates of response amplitude. The GMM acts as a signal filter ignoring spurious fluctuations in the firing rate.

### Isolating responses to perisaccadic probes

To isolate responses to the probe on trials with coincident saccades (R_Probe(Peri)_), we developed an algorithm to remove the inferred response to the saccade (R_Saccade(Shifted)_) from the observed combined response to the saccade and probe (R_Probe, Saccade_). R_Probe, Saccade_ was calculated as the mean firing rate in the window −200 to 500 ms from probe onset. To compute R_Saccade(Shifted)_, for each trial in a given set of perisaccadic probes, we time-shifted the mean response to saccades that occurred without any nearby probe (R_Saccade_) by the latency to the nearest saccade. For example, if the closest saccade to a given probe occurred 100 ms after a probe, we would take R_Saccade_ centered on the saccade onset and shift it forward in time by 100 ms. If the saccade happened –100 ms before the probe, we would shift R_Saccade_ backward in time by 100 ms. This latency-shifting procedure functions to place a typical response to saccades into a probe-relative time space. After computing these trial-by-trial latency-shifted forms of R_Saccade_, we average over trials to obtain R_Saccade(Shifted)_, an estimate of saccade-related activity relative to the onset of the probe for a given set of trials. Finally, we isolated the average perisaccadic visual response (R_Probe(Peri)_) by subtracting R_Saccade(Shifted)_ from R_Probe, Saccade_.

### Modulation index

To quantify saccadic modulation of visual responses, we formulated a modulation index (MI) that measures the magnitude difference between R_Probe(Extra)_ and R_Probe(Peri)_. First, we took the GMM fit to R_Probe(Extra)_ and refit it to R_Probe(Peri)_. We constrained the curve-fitting process such that the amplitude parameters were the only parameters we allowed to vary; latency and peak width parameters from the original fit were fixed. After refitting, we compared the amplitude of the largest component in the GMM fit to R_Probe(Extra)_ to the same component in the GMM refit to R_Probe(Peri)_. The difference of amplitudes was normalized to the amplitude of the largest component in R_Probe(Extra)_ such that the resulting MI reflects the loss or gain of amplitude as a fraction of the amplitude of the response to the extrasaccadic probe. For negative units, we inverted the sign of the responses so that changes in response amplitude were consistent with positively signed responses.

### Hypothesis testing with bootstrapping

To determine the statistical significance of modulation estimates, we developed a hypothesis test with bootstrapping that indicates the probability of observing a more extreme value of MI by chance. For each unit, we generated a null distribution of MI that represents the variability of MI under conditions in which we are certain the response is unmodulated. First, we down-sampled R_Probe(Extra)_ using 5% of extrasaccadic probes (the observed percentage of trials with S-P latency between 0 and 100 ms) selected randomly without replacement (R_Probe(Extra)_’). We then re-computed MI substituting R_Probe(Peri)_ with R_Probe(Extra)_’ such that whatever modulation is indicated can be exclusively attributed to sampling error. We repeated this procedure 1000 times and then compared the original MI to this null distribution of MI. A 2-tailed p-value was computed by dividing the frequency of more extreme values of MI by the size of the null distribution. An α level of 0.05 was used to classify saccadic modulation as significant.

### Neural population analysis

To identify how saccades modulate neural population dynamics, we measured the distance in N-dimensional space (N=number of units) between perisaccadic and extrasaccadic trajectories. For each population of simultaneously recorded visual SC units, we computed an N units x M time bins extrasaccadic population matrix based on R_Probe(Extra)_ for each unit in 10 ms time bins from −200 ms to 500 ms relative to the probe. We computed the perisaccadic population matrix similarly using R_Probe(Peri)_ for each unit. To analyze the distance between the extra and perisaccadic population activity, we found the largest euclidean distance between population vectors between 20 ms and 160 ms window relative to the probe. The distance was calculated independently for each 10 ms time bin and over the full range of the 10 S-P latencies (−500 to 500 ms in 100 ms latency bins). The distance was calculated by randomly drawing the number of perisaccadic events from the available extrasaccadic events (typically, the number of extrasacaddic probes exceeded the number of perisaccadic events by a factor of 100). In many sessions, the R_Probe(Peri)_ - R_Probe(Extra)_ distance was larger than the distances observed by bootstrapping. Thus, we approximated the bootstrapped distribution with a normal distribution and used a p-value of p < 0.001 to determine individual session significance. The p-values were corrected for multiple comparisons using Bonferroni correction.

### Quantification and statistical analysis

We used a Χ^2^ goodness of fit to determine if a sample of a categorical variable is uniformly distributed or distributed differently than another sample. We used Spearman’s Rank correlation or Pearson’s correlation to assess the correlation between ordinal variables. We used the Wilcoxon signed rank test to determine if a sample’s median varied significantly from 0. Bootstrapping was used to compute p-values associated with each MI.

## REFERENCES

1. Schroeder, C.E., Wilson, D.A., Radman, T., Scharfman, H., and Lakatos, P. (2010). Dynamics of Active Sensing and perceptual selection. Curr Opin Neurobiol 20, 172–176. 10.1016/j.conb.2010.02.010.

2. Hubel, D.H., and Wiesel, T.N. (1962). Receptive fields, binocular interaction and functional architecture in the cat’s visual cortex. J Physiol 160, 106–154.2.

3. Felleman, D.J., and Van Essen, D.C. (1991). Distributed hierarchical processing in the primate cerebral cortex. Cereb Cortex 1, 1–47. 10.1093/cercor/1.1.1-a.

4. Werblin, F.S. (2011). The retinal hypercircuit: a repeating synaptic interactive motif underlying visual function. J Physiol 589, 3691–3702. 10.1113/jphysiol.2011.210617.

5. Seabrook, T.A., Burbridge, T.J., Crair, M.C., and Huberman, A.D. (2017). Architecture, Function, and Assembly of the Mouse Visual System. Annu Rev Neurosci 40, 499–538. 10.1146/annurev-neuro-071714-033842.

6. Siegle, J.H., Jia, X., Durand, S., Gale, S., Bennett, C., Graddis, N., Heller, G., Ramirez, T.K., Choi, H., Luviano, J.A., et al. (2021). Survey of spiking in the mouse visual system reveals functional hierarchy. Nature 592, 86–92. 10.1038/s41586-020-03171-x.

7. Land, M.F. (1999). Motion and vision: why animals move their eyes. Journal of Comparative Physiology A: Sensory, Neural, and Behavioral Physiology 185, 341–352. 10.1007/s003590050393.

8. Zuber, B.L., and Stark, L. (1966). Saccadic suppression: elevation of visual threshold associated with saccadic eye movements. Exp Neurol 16, 65–79. 10.1016/0014-4886(66)90087-2.

9. Beeler, G.W. (1967). Visual threshold changes resulting from spontaneous saccadic eye movements. Vision Res 7, 769–775. 10.1016/0042-6989(67)90039-9.

10. Burr, D.C., Morrone, M.C., and Ross, J. (1994). Selective suppression of the magnocellular visual pathway during saccadic eye movements. Nature 371, 511–513. 10.1038/371511a0.

11. Diamond, M.R., Ross, J., and Morrone, M.C. (2000). Extraretinal Control of Saccadic Suppression. J. Neurosci. 20, 3449–3455. 10.1523/JNEUROSCI.20-09-03449.2000.

12. Wurtz, R.H. (1968). Visual cortex neurons: response to stimuli during rapid eye movements. Science 162, 1148–1150. 10.1126/science.162.3858.1148.

13. Robinson, D.L., and Wurtz, R.H. (1976). Use of an extraretinal signal by monkey superior colliculus neurons to distinguish real from self-induced stimulus movement. J Neurophysiol 39, 852–870. 10.1152/jn.1976.39.4.852.

14. Thiele, A., Henning, P., Kubischik, M., and Hoffmann, K.-P. (2002). Neural mechanisms of saccadic suppression. Science 295, 2460–2462. 10.1126/science.1068788.

15. Saul, A.B. (2010). Effects of fixational saccades on response timing in macaque lateral geniculate nucleus. Vis Neurosci 27, 171–181. 10.1017/S0952523810000258.

16. Krock, R.M., and Moore, T. (2016). Visual sensitivity of frontal eye field neurons during the preparation of saccadic eye movements. J Neurophysiol 116, 2882–2891. 10.1152/jn.01140.2015.

17. Ibbotson, M., and Krekelberg, B. (2011). Visual Perception and Saccadic Eye Movements. Curr Opin Neurobiol 21, 553–558. 10.1016/j.conb.2011.05.012.

18. Land, M.F., and Tatler, B.W. (2009). The human eye movement repertoire. In Looking and Acting: Vision and eye movements in natural behaviour, M. Land and B. Tatler, eds. (Oxford University Press), p. 0. 10.1093/acprof:oso/9780198570943.003.0002.

19. Meyer, A.F., O’Keefe, J., and Poort, J. (2020). Two Distinct Types of Eye-Head Coupling in Freely Moving Mice. Curr Biol 30, 2116–2130.e6. 10.1016/j.cub.2020.04.042.

20. Michaiel, A.M., Abe, E.T., and Niell, C.M. (2020). Dynamics of gaze control during prey capture in freely moving mice. eLife 9, e57458. 10.7554/eLife.57458.

21. Ambrad Giovannetti, E., and Rancz, E. (2024). Behind mouse eyes: The function and control of eye movements in mice. Neuroscience & Biobehavioral Reviews 161, 105671. 10.1016/j.neubiorev.2024.105671.

22. Skyberg, R.J., and Niell, C.M. (2024). Natural visual behavior and active sensing in the mouse. Current Opinion in Neurobiology 86, 102882. 10.1016/j.conb.2024.102882.

23. Parker, P.R.L., Martins, D.M., Leonard, E.S.P., Casey, N.M., Sharp, S.L., Abe, E.T.T., Smear, M.C., Yates, J.L., Mitchell, J.F., and Niell, C.M. (2023). A dynamic sequence of visual processing initiated by gaze shifts. Nat Neurosci 26, 2192–2202. 10.1038/s41593-023-01481-7.

24. Yonehara, K., Fiscella, M., Drinnenberg, A., Esposti, F., Trenholm, S., Krol, J., Franke, F., Scherf, B.G., Kusnyerik, A., Müller, J., et al. (2016). Congenital Nystagmus Gene FRMD7 Is Necessary for Establishing a Neuronal Circuit Asymmetry for Direction Selectivity. Neuron 89, 177–193. 10.1016/j.neuron.2015.11.032.

25. Gale, S.D., and Murphy, G.J. (2014). Distinct representation and distribution of visual information by specific cell types in mouse superficial superior colliculus. J Neurosci 34, 13458–13471. 10.1523/JNEUROSCI.2768-14.2014.

26. Li, Y., and Meister, M. (2023). Functional cell types in the mouse superior colliculus. eLife 12, e82367. 10.7554/eLife.82367.

27. Basso, M.A., and May, P.J. (2017). Circuits for Action and Cognition: A View from the Superior Colliculus. Annu Rev Vis Sci 3, 197–226. 10.1146/annurev-vision-102016-061234.

28. Oliveira, A.F., and Yonehara, K. (2018). The Mouse Superior Colliculus as a Model System for Investigating Cell Type-Based Mechanisms of Visual Motor Transformation. Front Neural Circuits 12, 59. 10.3389/fncir.2018.00059.

29. Basso, M.A., Bickford, M.E., and Cang, J. (2021). Unraveling circuits of visual perception and cognition through the superior colliculus. Neuron 109, 918–937. 10.1016/j.neuron.2021.01.013.

30. Isa, T., Marquez-Legorreta, E., Grillner, S., and Scott, E.K. (2021). The tectum/superior colliculus as the vertebrate solution for spatial sensory integration and action. Curr Biol 31, R741–R762. 10.1016/j.cub.2021.04.001.

31. Hoy, J.L., Yavorska, I., Wehr, M., and Niell, C.M. (2016). Vision Drives Accurate Approach Behavior during Prey Capture in Laboratory Mice. Curr Biol 26, 3046–3052. 10.1016/j.cub.2016.09.009.

32. Hoy, J.L., Bishop, H.I., and Niell, C.M. (2019). Defined Cell Types in Superior Colliculus Make Distinct Contributions to Prey Capture Behavior in the Mouse. Curr Biol 29, 4130–4138.e5. 10.1016/j.cub.2019.10.017.

33. Shang, C., Liu, Z., Chen, Z., Shi, Y., Wang, Q., Liu, S., Li, D., and Cao, P. (2015). BRAIN CIRCUITS. A parvalbumin-positive excitatory visual pathway to trigger fear responses in mice. Science 348, 1472–1477. 10.1126/science.aaa8694.

34. Shang, C., Chen, Z., Liu, A., Li, Y., Zhang, J., Qu, B., Yan, F., Zhang, Y., Liu, W., Liu, Z., et al. (2018). Divergent midbrain circuits orchestrate escape and freezing responses to looming stimuli in mice. Nat Commun 9, 1232. 10.1038/s41467-018-03580-7.

35. Ellis, E.M., Gauvain, G., Sivyer, B., and Murphy, G.J. (2016). Shared and distinct retinal input to the mouse superior colliculus and dorsal lateral geniculate nucleus. J Neurophysiol 116, 602–610. 10.1152/jn.00227.2016.

36. Gandhi, N.J., and Katnani, H.A. (2011). Motor functions of the superior colliculus. Annu Rev Neurosci 34, 205–231. 10.1146/annurev-neuro-061010-113728.

37. González-Rueda, A., Jensen, K., Noormandipour, M., de Malmazet, D., Wilson, J., Ciabatti, E., Kim, J., Williams, E., Poort, J., Hennequin, G., et al. (2024). Kinetic features dictate sensorimotor alignment in the superior colliculus. Nature 631, 378–385. 10.1038/s41586-024-07619-2.

38. Lee, P.H., Sooksawate, T., Yanagawa, Y., Isa, K., Isa, T., and Hall, W.C. (2007). Identity of a pathway for saccadic suppression. Proc Natl Acad Sci U S A 104, 6824–6827. 10.1073/pnas.0701934104.

39. Isa, T., and Hall, W.C. (2009). Exploring the Superior Colliculus In Vitro. J Neurophysiol 102, 2581–2593. 10.1152/jn.00498.2009.

40. Phongphanphanee, P., Mizuno, F., Lee, P.H., Yanagawa, Y., Isa, T., and Hall, W.C. (2011). A circuit model for saccadic suppression in the superior colliculus. J Neurosci 31, 1949–1954. 10.1523/JNEUROSCI.2305-10.2011.

41. Idrees, S., Baumann, M.P., Franke, F., Münch, T.A., and Hafed, Z.M. (2020). Perceptual saccadic suppression starts in the retina. Nat Commun 11, 1977. 10.1038/s41467-020-15890-w.

42. Idrees, S., Baumann, M.-P., Korympidou, M.M., Schubert, T., Kling, A., Franke, K., Hafed, Z.M., Franke, F., and Münch, T.A. (2022). Suppression without inhibition: how retinal computation contributes to saccadic suppression. Commun Biol 5, 692. 10.1038/s42003-022-03526-2.

43. Cafaro, J., Zylberberg, J., and Field, G.D. (2020). Global Motion Processing by Populations of Direction-Selective Retinal Ganglion Cells. J. Neurosci. 40, 5807–5819. 10.1523/JNEUROSCI.0564-20.2020.

44. Krishnamoorthy, V., Weick, M., and Gollisch, T. Sensitivity to image recurrence across eye-movement-like image transitions through local serial inhibition in the retina. eLife 6, e22431. 10.7554/eLife.22431.

45. Mani, A., Yang, X., Zhao, T.A., Leyrer, M.L., Schreck, D., and Berson, D.M. (2023). A circuit suppressing retinal drive to the optokinetic system during fast image motion. Nat Commun 14, 5142. 10.1038/s41467-023-40527-z.

46. Roska, B., and Werblin, F. (2003). Rapid global shifts in natural scenes block spiking in specific ganglion cell types. Nat Neurosci 6, 600–608. 10.1038/nn1061.

47. Sivyer, B., Tomlinson, A., and Taylor, W.R. (2019). Simulated Saccadic Stimuli Suppress ON-Type Direction-Selective Retinal Ganglion Cells via Glycinergic Inhibition. J Neurosci 39, 4312–4322. 10.1523/JNEUROSCI.3066-18.2019.

48. Summers, M.T., and Feller, M.B. (2022). Distinct inhibitory pathways control velocity and directional tuning in the mouse retina. Curr Biol 32, 2130–2143.e3. 10.1016/j.cub.2022.03.054.

49. Tien, N.-W., Pearson, J.T., Heller, C.R., Demas, J., and Kerschensteiner, D. (2015). Genetically Identified Suppressed-by-Contrast Retinal Ganglion Cells Reliably Signal Self-Generated Visual Stimuli. J Neurosci 35, 10815–10820. 10.1523/JNEUROSCI.1521-15.2015.

50. Kretschmer, F., Sajgo, S., Kretschmer, V., and Badea, T.C. (2015). A system to measure the Optokinetic and Optomotor response in mice. J Neurosci Methods 256, 91–105. 10.1016/j.jneumeth.2015.08.007.

51. Mathis, A., Mamidanna, P., Cury, K.M., Abe, T., Murthy, V.N., Mathis, M.W., and Bethge, M. (2018). DeepLabCut: markerless pose estimation of user-defined body parts with deep learning. Nat Neurosci 21, 1281–1289. 10.1038/s41593-018-0209-y.

52. Sibille, J., Gehr, C., Benichov, J.I., Balasubramanian, H., Teh, K.L., Lupashina, T., Vallentin, D., and Kremkow, J. (2022). High-density electrode recordings reveal strong and specific connections between retinal ganglion cells and midbrain neurons. Nat Commun 13, 5218. 10.1038/s41467-022-32775-2.

53. Montijn, J.S., Seignette, K., Howlett, M.H., Cazemier, J.L., Kamermans, M., Levelt, C.N., and Heimel, J.A. A parameter-free statistical test for neuronal responsiveness. eLife 10, e71969. 10.7554/eLife.71969.

54. Hafed, Z.M., and Krauzlis, R.J. (2010). Microsaccadic suppression of visual bursts in the primate superior colliculus. J Neurosci 30, 9542–9547. 10.1523/JNEUROSCI.1137-10.2010.

55. Chen, C.-Y., Ignashchenkova, A., Thier, P., and Hafed, Z.M. (2015). Neuronal Response Gain Enhancement prior to Microsaccades. Curr Biol 25, 2065–2074. 10.1016/j.cub.2015.06.022.

56. Chen, C.-Y., and Hafed, Z.M. (2017). A neural locus for spatial-frequency specific saccadic suppression in visual-motor neurons of the primate superior colliculus. J Neurophysiol 117, 1657–1673. 10.1152/jn.00911.2016.

57. Berman, R.A., Cavanaugh, J., McAlonan, K., and Wurtz, R.H. (2017). A circuit for saccadic suppression in the primate brain. J Neurophysiol 117, 1720–1735. 10.1152/jn.00679.2016.

58. Wang, L., Sarnaik, R., Rangarajan, K., Liu, X., and Cang, J. (2010). Visual receptive field properties of neurons in the superficial superior colliculus of the mouse. J Neurosci 30, 16573–16584. 10.1523/JNEUROSCI.3305-10.2010.

59. Wurtz, R.H. (2008). Neuronal mechanisms of visual stability. Vision Res 48, 2070–2089. 10.1016/j.visres.2008.03.021.

60. Jampel, R.S., and Shi, D.X. (1992). The primary position of the eyes, the resetting saccade, and the transverse visual head plane. Head movements around the cervical joints. Invest Ophthalmol Vis Sci 33, 2501–2510.

61. Stahl, J.S. (1999). Amplitude of human head movements associated with horizontal saccades. Exp Brain Res 126, 41–54. 10.1007/s002210050715.

62. Stahl, J.S. (2001). Eye-head coordination and the variation of eye-movement accuracy with orbital eccentricity. Exp Brain Res 136, 200–210. 10.1007/s002210000593.

63. Foulsham, T., Walker, E., and Kingstone, A. (2011). The where, what and when of gaze allocation in the lab and the natural environment. Vision Res 51, 1920–1931. 10.1016/j.visres.2011.07.002.

64. Ioannidou, F., Hermens, F., and Hodgson, T.L. (2016). The Central Bias in Day-to-Day Viewing. Journal of Eye Movement Research 9. 10.16910/jemr.9.6.6.

65. Burlingham, C.S., Sendhilnathan, N., Komogortsev, O., Murdison, T.S., and Proulx, M.J. (2024). Motor “laziness” constrains fixation selection in real-world tasks. Proc Natl Acad Sci U S A 121, e2302239121. 10.1073/pnas.2302239121.

66. Hoggarth, A., McLaughlin, A.J., Ronellenfitch, K., Trenholm, S., Vasandani, R., Sethuramanujam, S., Schwab, D., Briggman, K.L., and Awatramani, G.B. (2015). Specific wiring of distinct amacrine cells in the directionally selective retinal circuit permits independent coding of direction and size. Neuron 86, 276–291. 10.1016/j.neuron.2015.02.035.

67. Huang, X., Rangel, M., Briggman, K.L., and Wei, W. (2019). Neural mechanisms of contextual modulation in the retinal direction selective circuit. Nat Commun 10, 2431. 10.1038/s41467-019-10268-z.

68. Matsumoto, A., Agbariah, W., Nolte, S.S., Andrawos, R., Levi, H., Sabbah, S., and Yonehara, K. (2021). Direction selectivity in retinal bipolar cell axon terminals. Neuron 109, 2928–2942.e8. 10.1016/j.neuron.2021.07.008.

69. Ghitani, N., Bayguinov, P.O., Vokoun, C.R., McMahon, S., Jackson, M.B., and Basso, M.A. (2014). Excitatory Synaptic Feedback from the Motor Layer to the Sensory Layers of the Superior Colliculus. J Neurosci 34, 6822–6833. 10.1523/JNEUROSCI.3137-13.2014.

70. Marino, A.C., and Mazer, J.A. (2016). Perisaccadic Updating of Visual Representations and Attentional States: Linking Behavior and Neurophysiology. Front Syst Neurosci 10, 3. 10.3389/fnsys.2016.00003.

71. Miura, S.K., and Scanziani, M. (2022). Distinguishing externally from saccade-induced motion in visual cortex. Nature 610, 135–142. 10.1038/s41586-022-05196-w.

72. Essig, J., Hunt, J.B., and Felsen, G. (2021). Inhibitory neurons in the superior colliculus mediate selection of spatially-directed movements. Commun Biol 4, 719. 10.1038/s42003-021-02248-1.

73. Arnold Jon Low-Friction Rodent-Driven Belt Treadmill. Janelia Research Campus. https://www.janelia.org/open-science/low-friction-rodent-driven-belt-treadmill.

74. De Franceschi, G., and Solomon, S.G. (2018). Visual response properties of neurons in the superficial layers of the superior colliculus of awake mouse. The Journal of Physiology 596, 6307–6332. 10.1113/JP276964.

